# Autoantibodies in patients with arrhythmogenic cardiomyopathy activate GSK-3β resulting in a loss of cardiomyocyte cohesion

**DOI:** 10.1101/2025.06.25.661311

**Authors:** Soumyata Pathak, Konstanze Stangner, Ellen Kempf, Sina Moztarzadeh, Matthias Hiermaier, Marlene Rauschmayer, Tatjana Williams, Andreas Stengl, Brenda Gerull, Ruth Biller, Sebastian Clauss, Stefan Kääb, Tomo Šarić, Sunil Yeruva, Jens Waschke

**Author notes:** These authors contributed equally (Sunil Yeruva, Jens Waschke). **Corresponding authors:** Sunil Yeruva and Jens Waschke, Chair of Vegetative Anatomy, Institute of Anatomy, Ludwig-Maximilians-University, Pettenkoferstrasse 11, 80336 Munich, Germany; and. Conflict of interest: The authors have declared that no conflict of interest exists.

## Abstract

Arrhythmogenic cardiomyopathy (ACM) is an inherited cardiac desmosome disease, as more than 50% of affected patients carry pathogenic variants in desmosome protein-coding genes., In this study, we focused on the role and mechanisms of pathogenic and non-pathogenic autoantibodies against intercalated disc (ICD) proteins such as desmoglein2 (DSG2) in ACM patients, healthy relatives and murine ACM models. IgG fractions from ACM patients and healthy relatives, but not murine ACM model-derived or healthy control IgGs, revealed positive ICD staining. Antibodies reducing the loss of cardiomyocyte cohesion were found in three out of five ACM patients. Pathogenic autoantibodies, bound to DSG2 in hiPSC-CMs, cleaved DSG2 and reduced DSG2 interaction at the molecular level. We investigated GSK-3β contribution to the loss of cardiomyocyte cohesion and observed that GSK-3β reduced baseline cardiomyocyte cohesion in cultured cardiomyocytes and cardiac slices. Pathogenic ACM-IgGs activated GSK-3β upstream of p38MAPK, leading to phosphorylation and junctional loss of β-catenin. GSK-3β inhibition rescued the loss of cell cohesion induced by ACM-IgGs in ACM hiPSC-CMs. Pathogenic autoantibodies targeting DSG2 are present in ACM patients and impair cardiomyocyte cohesion in a GSK-3β-dependent manner. In contrast, autoantibodies are absent in murine ACM models and are non-pathogenic in some patients, healthy relatives.

## Introduction

Cardiomyocytes within the functional syncytium of the heart are electromechanically connected via intercalated discs (ICD) that comprise desmosomes and adherens junction proteins, which provide adhesive contact to adjacent cardiomyocytes, interact with the cytoskeleton, and participate in crosstalk with gap junctions and other downstream proteins (1, 2). The desmosomal proteins: desmoplakin (DSP), plakophilin 2 (PKP2), desmoglein 2 (DSG2), plakoglobin (JUP), and desmocollin 2 (DSC2) are encoded by genes that undergo pathogenic variations, with varying prevalence, in more than 50% of individuals with arrhythmogenic cardiomyopathy (ACM) (1, 3-8). Nevertheless, pathogenic variations found in ACM patients are not confined solely to desmosomal proteins but rather are rarely observed in non-desmosomal proteins such as N-cadherin (N-CAD), transmembrane protein 43 (TMEM 43), phospholamban (PLN), ryanodine receptor 2 (RYR 2), and Titin (TTN) (9-11).

ACM is a cause of sudden cardiac death (SCD) in young individuals, especially athletes and in some patients, SCD could be the first presentation of the disease (12-14). Clinically, ACM can be divided into four phases: concealed phase with no apparent symptoms, an overt phase where palpitations and arrhythmias occur along with some overt structural alterations, and an end phase where structural and electrical abnormalities result in ventricular dilation and the fourth phase leads to dilated cardiomyopathy and biventricular failure (12). Although genetic determinants have been identified, 35-50% of patients are mutation negative, indicating complex etiologies of myocardium replacement to fibrofatty tissue resulting in aneurysms and wall weakening that impairs electrical conduction, inflammation and cardiac failure involving non-genetic anomalies (12, 15).

Therefore, it was highly interesting that autoantibodies against DSG2 were detected in ACM patients (16), which were found to bind to ICDs (17) and in addition to genetic changes, could further contribute to ACM pathogenesis. This is possible because in pemphigus, a desmosome disease of the skin, autoantibodies against DSG1 and DSG3 are well established to be the main factors contributing to the disease pathogenesis by impairing cell cohesion (18, 19) In line with that, a recent study demonstrated that autoantibodies derived from ACM patients have catalytic properties against DSG2 and N-CAD and disrupt cardiomyocyte cohesion by activating p38MAPK, a phenomenon observed for keratinocytes in pemphigus (20). However, very little is known about the role of autoantibodies in the pathogenesis of ACM. Recent studies, utilizing murine ACM models (21) and hiPSC-derived cardiomyocytes from ACM patients (22) provide evidence that desmosomal adhesion could also play a role in ACM pathogenesis (18). In this study, we found that autoantibodies were present in all ACM patients and healthy relatives but were absent in murine ACM models and healthy control individuals. However, autoantibodies were pathogenic in some ACM patients, reduced cardiomyocyte cohesion but were non-pathogenic in HRs indicating that autoantibodies develop independently of the disease-causing pathogenic variant, yet with a familial preference, suggesting underlying genetic predisposition. Pathogenic autoantibodies bound to DSG2 in hiPSC-CMs and interfered with DSG2 binding on the molecular level. Finally, we identified that pathogenic autoantibodies reduced cardiomyocyte cohesion in a GSK-3β-dependent manner in ACM patient-derived hiPSC-CMs. Our data reveal that, in addition to pathogenic variants, autoantibodies could also contribute to the activation of GSK-3β observed in ACM patients and mouse models and may further aggravate ACM pathogenesis.

## Results

### ACM patient IgGs bind to DSG2 in hiPSC-CMs and cause loss of cohesion

Previous studies have established the formation of autoantibodies against DSG2 (16) or intercalated disc proteins in an ACM patient cohort (23). In a recent study, we established that certain patients with ACM produce catalytic antibodies against DSG2 and N-CAD, resulting in a loss of cell cohesion in murine cardiomyocytes (20). Here, we investigated whether this loss of cell cohesion also occurs in human cardiomyocytes using dissociation assays with cardiomyocytes derived from a healthy male (NP0040-8). We treated human induced pluripotent stem cell-derived cardiomyocytes (hiPSC-CMs) with immunoglobulin G (IgG) from ACM patients ACM 1 and ACM 15, as previously studied (24). After 24 h of incubation, the detached monolayers were shaken to apply mechanical force, and the resulting fragments were counted against those from control IgG (HC-IgG). The results indicated that hiPSC-CMs treated with IgGs from ACM 1 and ACM 15 generated significantly more fragments than those treated with HC-IgG, confirming that ACM patient IgGs can cause a loss of cohesion in human cardiomyocytes, similar to the effects observed in HL-1 cells, in our previous study **(Fig. 1A)**. We also observed that ACM 1-IgG but not IgG from two healthy controls (HC 1 and HC 2) detected DSG2-His recombinant protein in ELISA (**Fig. 1B)**. In contrast, an ELISA with increasing concentrations of DSG2-Fc (100 -1000 nM) resulted in non-specific binding of ACM 1-IgG as was observed with the Fc-tag control (Fc-tag Ctrl) ELISA. **(Supplementary Fig. 1A).** These results confirmed that, as previously observed, detecting anti-DSG2 antibodies using DSG2-Fc ELISA may lead to false-positive results.

**Figure 1.**
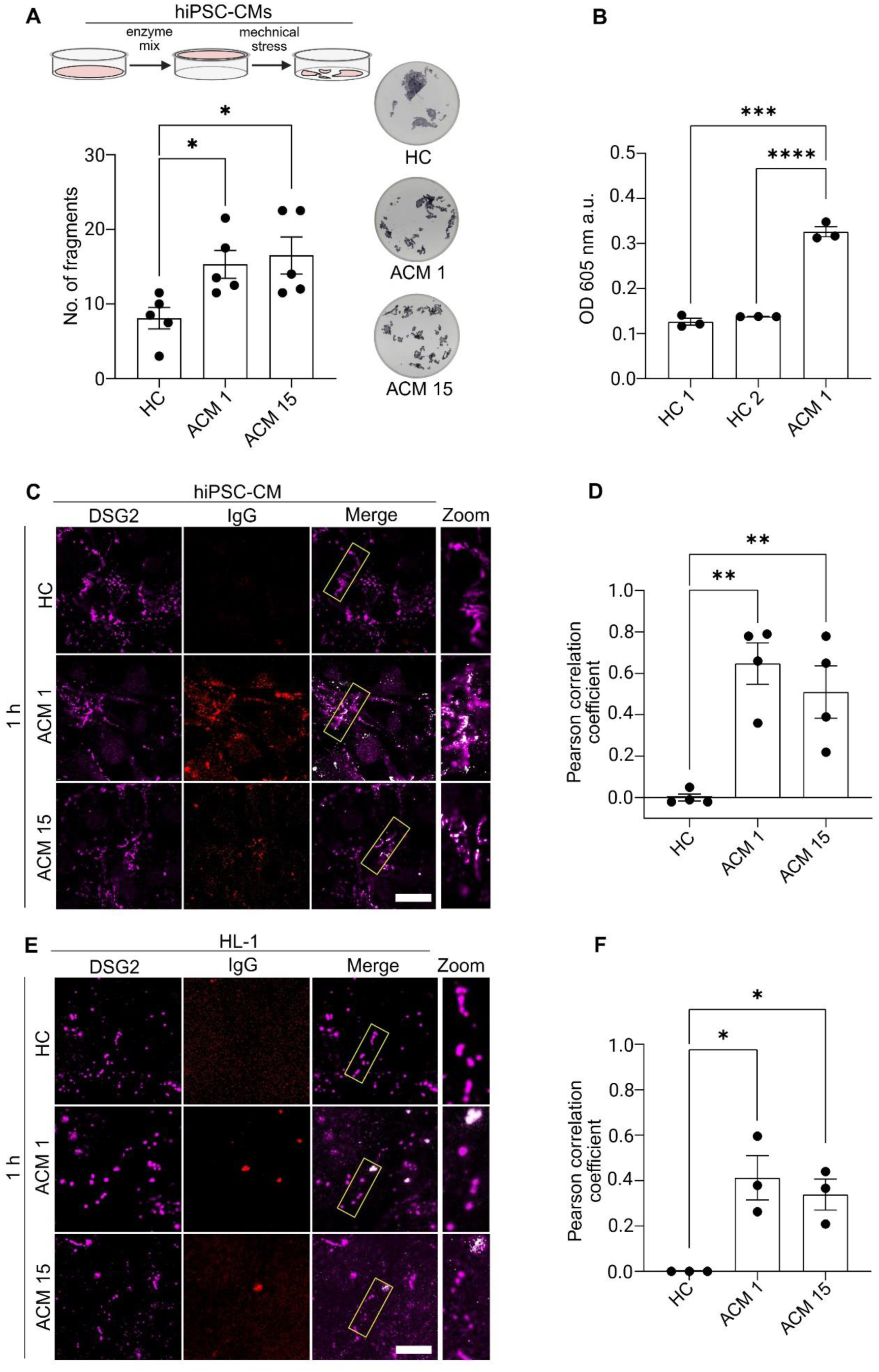
ACM-IgGs cause loss of cohesion and colocalize with DSG2 in hiPSC-CMs. **(A)** Dispase based dissociation assay was performed in hiPSC derived cardiomyocytes (hiPSC-CMs, Healthy male, NP0040-8) after 24 h incubation with respective ACM 1 and ACM 15-IgGs and grouped healthy Ctrl-IgG (HC). Compared to HC-IgG the increase in fragments in ACM-IgGs shows loss of cohesion. Representative images of cell fragments for each IgGs from 5 experimental repeats. Data are presented as mean ± SEM. One-way ANOVA with the Holm-Šidák multiple comparisons test was performed for statistical significance. **(B)** ELISA was performed using 500 nM/75 µg/ml concentration of IgGs from 2 healthy Ctrl (HC 1, HC 2) and ACM 1-IgG incubated for 1 h to detect extracellular DSG2-His tagged protein (2.5 µg/ml). Paired t-test was performed to compare ACM 1 to that of HC 1 and HC 2. Immunofluorescence staining was performed to determine the colocalization of anti-DSG2 antibodies in **(C)** hiPSC-CMs (Healthy male, NP0040-8) and **(E)** HL-1 cells (Murine atrial cardiomyocytes) after incubating with respective ACM 1, ACM 15 and HC-IgGs for 1 h. The zoom from merged images shows the colocalized ACM-IgGs with DSG2 compared to HC. Images represent analysis from 3 independent repeats (Scale bar 10 µm). **(D and F)** The colocalization area intensities were quantified using Pearson correlation coefficient analysis, where 0 is no correlation, and values between 0 and 1 are positive correlation. One-way ANOVA with the Holm-Šidák multiple comparisons test was performed for statistical significance. *p ≤ 0.05, **p ≤ 0.005, ***p ≤ 0.0005, ****p ≤ 0.00005.

Previous studies have demonstrated the presence of autoantibodies against DSG2 in fixed human cardiac tissue (16, 23, 24). Therefore, we tested whether IgGs from patients with ACM bind to DSG2 in living hiPSC-CMs and HL-1 cells. We incubated cardiomyocytes with the respective IgGs for 1 and 24 h and performed confocal and STED high-resolution imaging to detect the colocalization of IgGs with DSG2 at cell borders. After just 1 h incubation **(Fig. 1C and E),** ACM-IgGs colocalized with DSG2. Colocalization intensity of IgG with DSG2 was quantified using the Pearson correlation coefficient **(Fig. 1D and F).** Notably, after 24 h, we observed that IgGs had been internalized (data not shown). The secondary goat anti-human antibody control alone for both hiPSC-CMs and HL-1 cells was not cross-reactive with DSG2 **(Supplementary Fig. 1B and C).**

### New ACM patient cohort IgGs show anti-ICD antibodies which in part cause cardiomyocyte cohesion loss

We collected blood samples from five new ACM patients (three with pathogenic variants in PKP2, one in DSP, and one in the DSG2 gene **(Supplementary Table 1)**, along with three healthy relatives (HR), to further evaluate the pathogenicity of ACM patient IgGs. Therefore, we first established whether the new pool of IgG fractions from patients with different mutations causes a loss of cardiomyocyte cohesion. We performed dissociation assays with the new pool of ACM-IgGs and HR-IgGs in HL-1 cells. ACM 22, ACM 26 and ACM 27-IgGs induced loss of cardiomyocyte cohesion, whereas ACM 28, ACM 29 and HR-IgGs did not **(Fig. 2A and B)**.

**Figure 2.**
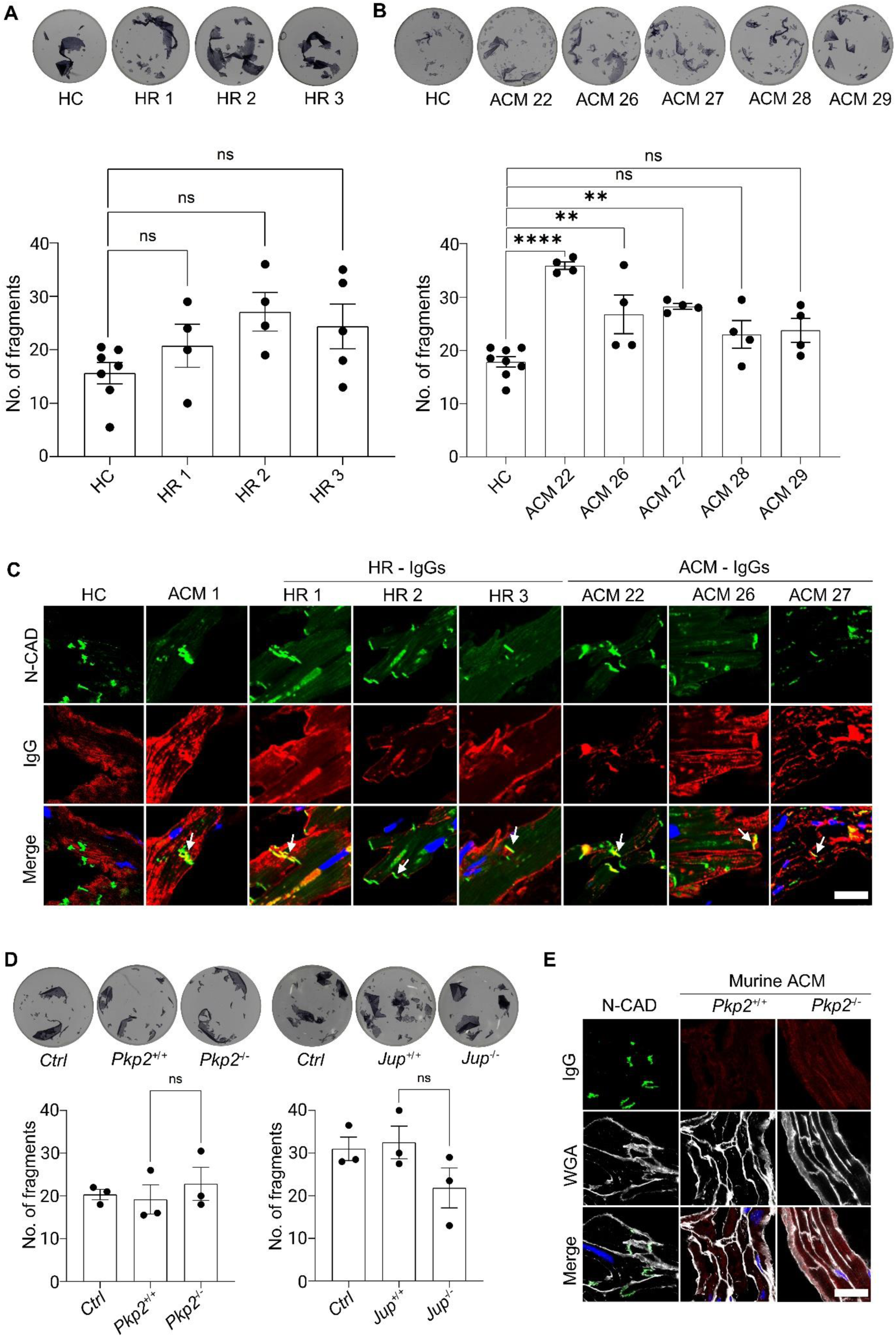
ACM-IgGs causing loss of cardiomyocyte cohesion are confined only to human. Dispase-based dissociation assay performed in HL-1 cells after 24 h incubation with respective (A) 3 HR-IgGs, HC-IgG, and **(B)** 5 ACM-IgGs. Representative images of cell fragments from 4 - 8 experimental repeats are displayed. Data are presented as mean ± SEM. One-way ANOVA with the Holm-Šidák multiple comparisons test was performed for statistical significance, **p ≤ 0.005, ***p ≤ 0.0005 and ns-not significant. **(C)** Immunostaining was performed using 3 HR-IgGs, 4 ACM-IgGs (ACM 1-IgG -positive control) and HC-IgG on human ventricular cardiac tissue. N-CAD was used as an ICD marker. The white arrow in the merge shows the overlap of all HR and ACM-IgGs with N-CAD except HC-IgG. Images represent 3 experimental repeats (Scale bar 10µm). **(D)** Dissociation assay in HL-1 cells was performed using *Pkp2^-/-^* and *Jup^-/-^* murine ACM-IgGs and their respective wild-type controls, after 24 h incubation. The wells represent cell fragments from 3 repeats. Data are presented as mean ± SEM. One-way ANOVA with the Holm-Šidák multiple comparisons test was performed to test statistical significance, ns-not significant. **(E)** Murine cardiac tissue used for immunostaining analysis to detect anti-ICD antibodies in murine ACM model. *Pkp2^-/-^* and *Pkp2^+/+^* -IgGs were incubated for 24 h and WGA was used as membrane marker. Image represents 3 experimental repeats (Scale bar 10 µm).

Here, we utilized a grouped control IgG (Grouped HC) prepared from four healthy donors as a cost-effective approach. However, to ensure that the observed effect with the grouped control IgGs does not mask individual IgG effects, we performed dissociation assays in HL-1 cells and immunostaining with human cardiac slices. We did not observe any increase in the number of fragments nor any staining of ICDs with individual IgG samples (HC 3, HC 4, HC 5, HC 6) compared to the grouped control IgG (Grouped HC) **(Supplementary Fig. 2A and B)**. Next, immunostaining was carried out with human ventricular cardiac tissue following incubation with the new pool of ACM-IgGs, three HR-IgGs, HC-IgG, and ACM 1 as a positive control. All ACM patients and HR-derived IgGs displayed positive staining of ICD, which overlapped with N-CAD, indicating the presence of anti-ICD antibodies **(Fig. 2C)** in all ACM-IgG and in HR-IgGs. Similarly, the goat anti-human antibody control on human cardiac tissue sections showed an intracellular staining, indicating mild cross-reactivity with human antigens **(Supplementary Fig. 2C).**

Because autoantibodies against ICD were present in the sera of ACM patients, we raised the question whether autoantibodies are present in murine models of ACM. To investigate this, we isolated blood from *Pkp2^-/-^* and *Jup^-/-^* ACM models and their respective wild-type littermates. IgGs were utilized for Immunostaining and dissociation assay. Incubation of HL-1 cells with the murine *Pkp2^-/-^* and *Jup^-/-^* IgG fractions for 24 h did not show reduced cohesion of cardiomocytes **(Fig. 2D)**. Immunostaining on murine ventricular cardiac tissue after incubation with IgGs did not stain ICD (WGA staining–membrane marker) **(Fig. 2E)** alongside the secondary goat anti-mouse antibody did not show cross reactivity on murine cardiac tissue **(Supplementary Fig. 2E).** These results demonstrate that autoantibodies targeting ICD proteins such as DSG2 and causing loss of cardiomyocyte cohesion are confined only to humans.

### ACM patients IgGs are catalytic and reduce homophilic DSG2 binding

Our previous study has shown that catalytic autoantibodies against DSG2 and N-CAD were present in the ACM patients (24). Therefore, the new pool of ACM-IgGs, which poses a strong reduction in cardiomyocyte cohesion, was also assessed for their catalytic behaviour against DSG2 protein using an in vitro cleavage assay. ACM-IgGs and HC-IgG were pre-incubated for 24 h with 200 ng of DSG2-Fc protein with protease inhibitor (PI) and without protease inhibitor (C) in cleavage assay buffer. The extracellular 100 kDa DSG2 band was detected in ACM 1 (positive control), ACM 26 and ACM 27-IgGs in the absence of PI (C). However, no cleaved DSG2 was detected in the presence of PI or after incubation with ACM 22-IgG **(Fig. 3A).** To measure the effect of ACM-IgGs on the relative binding frequency of DSG2, we utilized cell-free AFM experiments, in which a DSG2-Fc-coated cantilever was brought into contact with a DSG2-Fc-coated mica sheet and then retracted. The resulting force-distance-curves showed that after 1 h incubation the relative DSG2 binding frequency is reduced for all ACM-IgGs whereas it remains unchanged when incubating with HC-IgG **(Fig. 3B)**.

**Figure 3.**
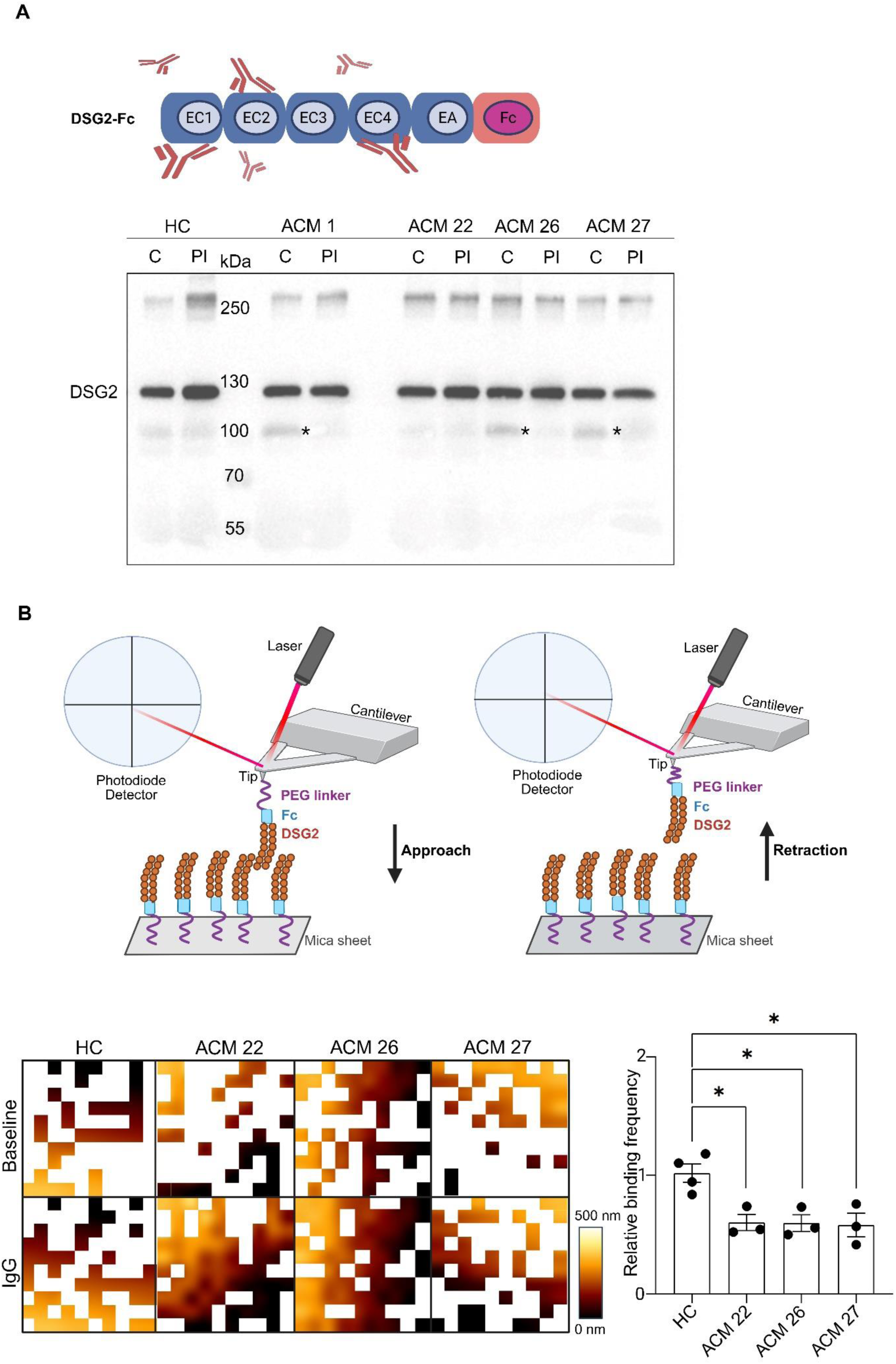
ACM patients IgGs are catalytic and reduce homophilic DSG2 binding. **(A**) In vitro DSG2-Fc cleavage assay was performed on ACM 22, ACM 26, ACM 27 and HC-IgGs in cleavage assay buffer for 24 h with protease inhibitor (PI) and without protease inhibitor (C). Western blots were carried out to detect the extracellular DSG2 band using an anti-DSG2 antibody that detects the extracellular domain of DSG2. Image blot represents image from 3 experimental repeats. * represents the cleaved DSG2 fragment. **(B)** The DSG2 homophilic binding in cell-free conditions was measured by AFM. Recombinant DSG2 tagged Fc fragments were attached to cantilever tip and mica sheet via PEG linker. The deflection of cantilever tip detected by laser and reflected on photodiode detector depicts an interaction event (Approach and retraction of DSG2-Fc) produced as Force-distance curve (FDC) along with adhesion maps (depicted below the cartoon) were generated by overlaying the position of occurring unbinding events with the mica topography maps. Each data point represents 1000 FDC after the 1 h incubation with respective ACM and HC-IgGs. The relative binding frequency is the resultant of FDC analyzed and normalized to baseline FDC measurements. Data represents as mean ± SEM of 3 - 4 experimental repeats. One-way ANOVA with Holm-Šidák multiple comparisons test was performed, *p<0.05.

### Autoantibodies in ACM patients cause GSK-3β activation

GSK-3β was found to be upregulated and present at the ICD in ACM patients (25). However, whether GSK-3β modulates cardiomyocyte cohesion and may contribute to ACM pathogenesis is unknown. To test activation of GSK-3β by ACM-IgGs, we performed Triton-X-100 extraction assays. The ACM-IgGs and HC were incubated for 1 and 24 h on HL-1 cells, followed by the separation of cytoskeletal from non-cytoskeletal proteins in Triton X-100 extraction buffer and Western blot. The respective no stain labelling of the membrane was used as loading control **(Supplementary Fig. 3A-C)** First, signalling analysis was performed after incubation of cells with ACM 1-IgG which revealed significant activation of GSK-3β, as indicated by reduced phosphorylation of GSK-3β at Ser 9, and activation of p38MAPK in the non-cytoskeleton bound fraction after 1 h **(Fig. 4D and F)**. Further, we observed an increase in phosphorylation of β-cat at Ser 33/37/Thr 41 compared to HC in the cytoskeleton-bound fraction, which may indicate that activated GSK-3β could phosphorylate β-cat bound to cell junctions. Next, we tested ACM 22, ACM 26 and ACM 27-IgG after 1 h **(Fig. 4C and E**). Consistently, ACM 26-IgG caused significant GSK-3β and p38MAPK activation as well as phosphorylation of β-cat. In contrast, ACM22-IgG exhibited a trend in activating GSK-3β and induced significant phosphorylation of β-cat and activation of p38MAPK. ACM 27-IgG significantly activated GSK-3β and showed a trend in p38MAPK activation and β-cat phosphorylation. GSK-3β and p38MAPK activation were absent after incubation for 24 h **(Supplementary Fig. 3D and E).** When normalized to HC, ACM-IgGs neither had any effects on protein levels of N-CAD and DSG2 after incubation for 24 h (**Fig. 4A and B)** nor on other signaling proteins **(Supplementary Fig. 3F and G).**

**Figure 4.**
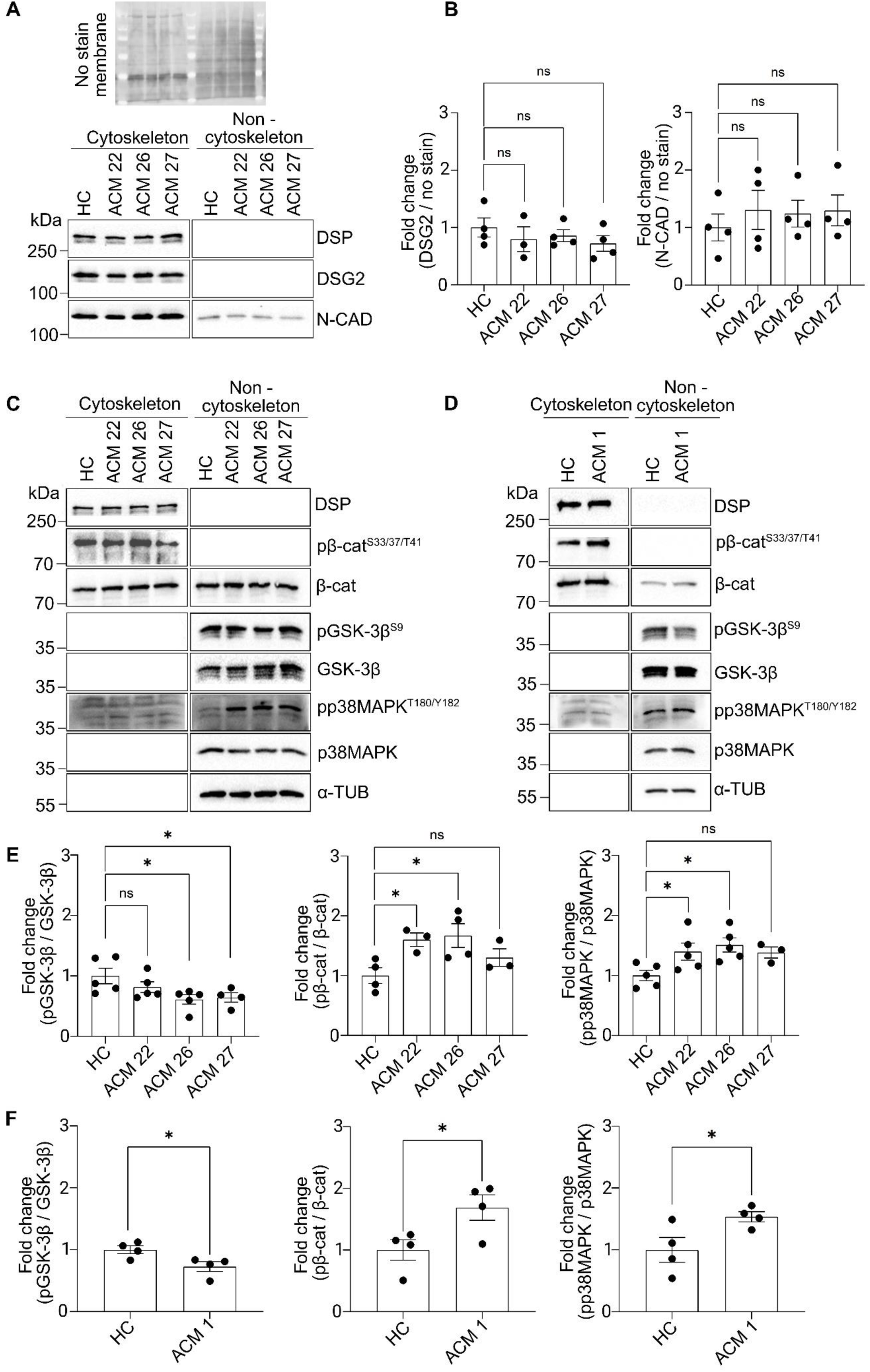
Autoantibodies in ACM patients cause GSK-3β activation. **(A)** Triton X-100 extraction assay was performed in HL-1 cells lysates after 24 h incubation with ACM 22, ACM 26, ACM 27 and HC-IgGs, to separate cytoskeleton and non-cytoskeleton proteins followed by Western blot for DSG2 and N-CAD protein analysis, No stain membrane was used as loading control. **(B)** Bar graphs represent the fold change of ratio between the DSG2 and N-CAD to loading control normalized to HC. **(C and D)** Western blots from Triton X-100 lysates of HL-1 cells after 1 h ACM-IgGs and HC-IgG treatments. **(E and F)** Quantification of Western blots from **(C and D)** are represented as fold change of ratio between phosphorylated protein to that of total protein (pGSK-3β / GSK-3β, pβ-cat / β-cat and pp38MAPK / p38MAPK) normalized to control group IgG (HC). Data are presented as mean ± SEM of 3 - 5 repeats. Statistical analysis was performed by one-way ANOVA with the Holm-Šidák multiple comparisons test and un-paired *t*-test, *p<0.05 and ns-not significant.

### ACM-IgGs cause loss of cohesion in hiPSC-CMs which was ameliorated by inhibition of GSK-3β

We further investigated the effects of ACM-IgGs in cardiomyocytes of ACM patients. We used hiPSCs from a 14-year-old female ACM patient who died of sudden cardiac death (NP0151-11F, index patient) and from her mother (NP0147-4, healthy relative). These hiPSCs characterization and differentiation into cardiomyocytes were described previously (22). The index patient carried a heterozygous pathogenic variant in DSP c.2854G>T, leading to a truncation of the desmoplakin protein at p. Glu952*, and presented with a biventricular phenotype, as well as confirmed ARVC in the family according to the Task Force Criteria of 2010 (15). Dissociation assays were performed on hiPSC-CMs from both healthy and ACM patients. Both healthy and ACM hiPSC-CMs were incubated with IgGs for 24 h **(Fig. 5A and 5B)**. Here, we chose ACM 1-IgG as a representative of our previous patient cohort and ACM 22-IgG because we observed a strong effect on the loss of cohesion in HL-1 cells. Both ACM-IgG fractions significantly increased the number of fragments in ACM hiPSC-CMs, whereas in healthy hiPSC-CMs, ACM 1-IgG but not ACM 22-IgG reduced cardiomyocyte cohesion.

**Figure 5.**
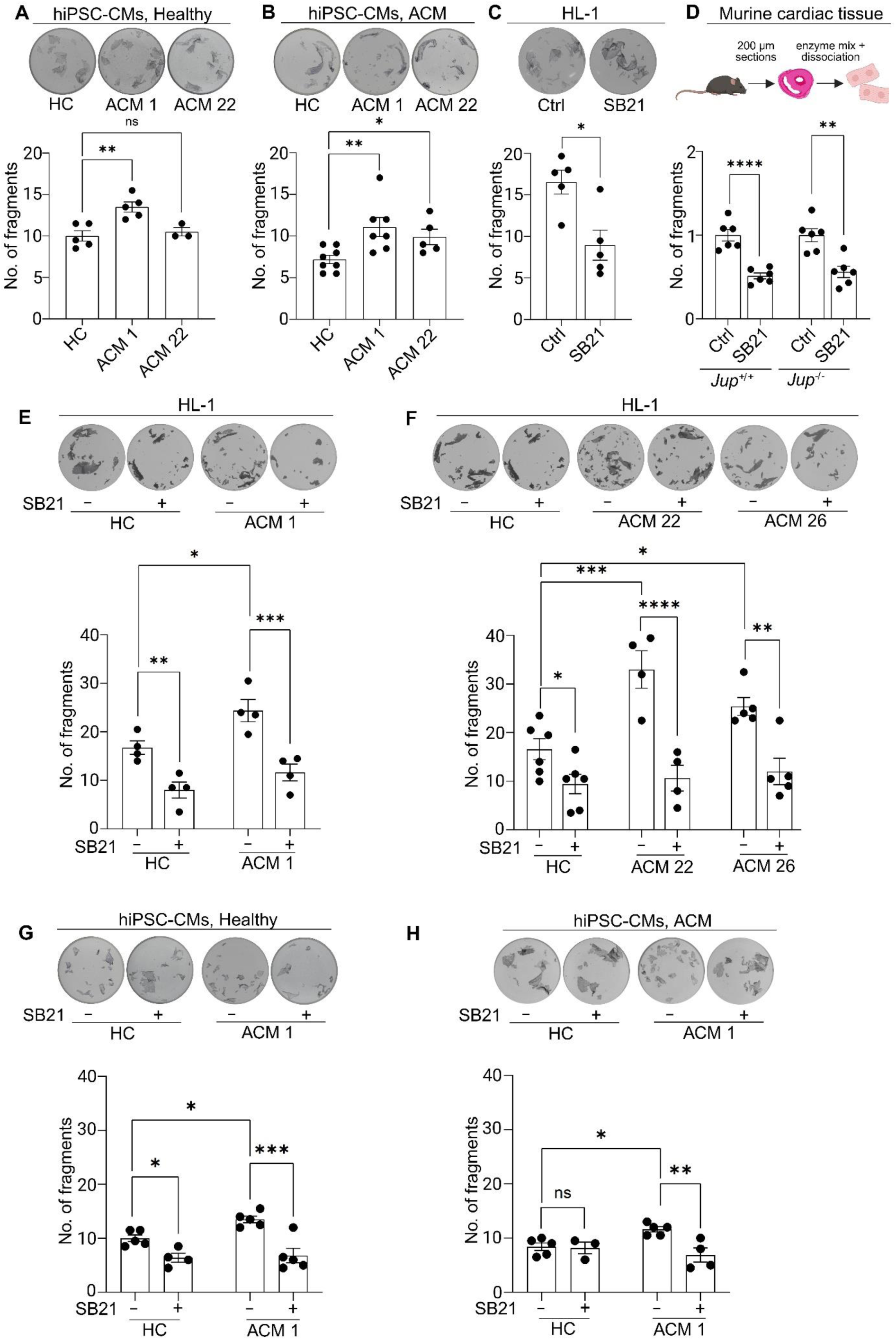
Inhibition of GSK-3β ameliorates loss of cohesion caused by ACM-IgGs in hiPSC-CMs. Dispase-based dissociation assay was performed in hiPSC-CMs from **(A)** healthy (NP147-4) and **(B)** ACM patients (NP151-11F) after 24 h incubation with ACM 1, ACM 22 and HC-IgGs. Representative images of cell fragments from 3 - 8 experimental repeats are displayed. Dissociation assay utilizing **(C)** HL-1cells and **(D)** murine cardiac tissue from *Jup^+/+^* and *Jup^-/-^* ACM model after 1 h incubation with SB21 was performed. Representative images of cell fragments from 5 - 6 repeats are displayed. The cardiac tissue dissociation of *Jup^-/-^* was normalized to *Jup^+/+^*(wild type). Data are presented as mean ± SEM. Statistical analysis was performed by one–way ANOVA with the Holm-Šidák multiple comparisons test for (A) and (B), un-paired *t*-test was performed to compare SB21 to Ctrl in (C) and two-way ANOVA with Holm-Šidák multiple comparisons was performed in (D). **(E and F)** HL-1 cells utilized to analyse the protective effect of SB21, on ACM-IgGs caused loss of cohesion, by dissociation after 24 h incubation of ACM and HC-IgGs fractions with (+) and without (-) SB21. The images represent the respective cell fragments from 4 - 6 repeats. Similarly, dissociation assays on hiPSC-CMs from **(G)** healthy and **(H)** ACM patients were performed after 24 h incubation with ACM 1 and HC-IgG with (+) and without (-) SB21. The images represent the respective cell fragments from 3 - 5 repeats. Data are presented as mean ± SEM. Statistical analysis was performed by two– way ANOVA with the Holm-Šidák multiple comparisons test. *p<0.05, **p<0.005, ***p<0.0005, ****p<0.00005 and ns-not significant.

First by using SB216763 (SB21), a small molecule GSK-3β inhibitor, which is known to rescue ACM pathogenesis in murine ACM models and hiPSC-CMs from ACM patients (26) (27). We tested whether inhibition of GSK-3β would affect baseline cardiomyocyte cohesion. We used a dissociation assay to evaluate the protective effect of SB21 in HL-1 cells and murine cardiac slices from wild-type mice and the *Jup^-/^*^-^ ACM model. Dissociation assays revealed a significant increase in the cohesion of HL-1 cells, wild-type, and *Jup^-/-^* cardiomyocytes after SB21 incubation **(Fig. 5C and D)**, indicating a negative role of GSK-3β in the regulation of cardiomyocyte cohesion. Then we incubated HL-1 cells with SB21 for 30 min before adding ACM 1, ACM 22, ACM26-IgG, and their respective controls for 24 h. SB21 significantly rescued ACM-IgG-mediated loss of cardiomyocyte cohesion **(Fig. 5E and F)**. Similarly, hiPSC-CMs from healthy and ACM patients were incubated with ACM 1-IgG, both with and without SB21. GSK-3β inhibition completely abolished the negative effects of ACM 1-IgG in healthy and ACM hiPSC-CMs **(Fig. 5G and H)**.

### Inhibiting GSK-3β restores β-cat along cell junctions and inactivates p38MAPK

Finally, we investigated the effect of GSK-3β inhibition on β-cat levels because phosphorylated β-cat is prone to proteasomal degradation. Triton X-100 assays were performed in the HL-1 cells after 1 h incubation with ACM 1 or ACM 26-IgGs with and without SB21 **(Fig. 6A-D)**. Western blot analysis revealed that ACM 1-IgG-mediated β-cat phosphorylation and p38MAPK activation were significantly ameliorated by SB21, suggesting that p38MAPK activation is GSK-3β dependent. After incubation with ACM 26-IgG, an increasing trend for β-cat phosphorylation and a significant decrease for p38MAPK phosphorylation were observed, when compared with HC. Immunostaining in HL-1 cells after 24 h incubation with ACM 1 and HC-IgGs revealed that ACM 1-IgG induced fragmentation and decrease of β-cat along cell junctions, which was rescued by SB21 **(Fig. 6E and F).**

**Figure 6.**
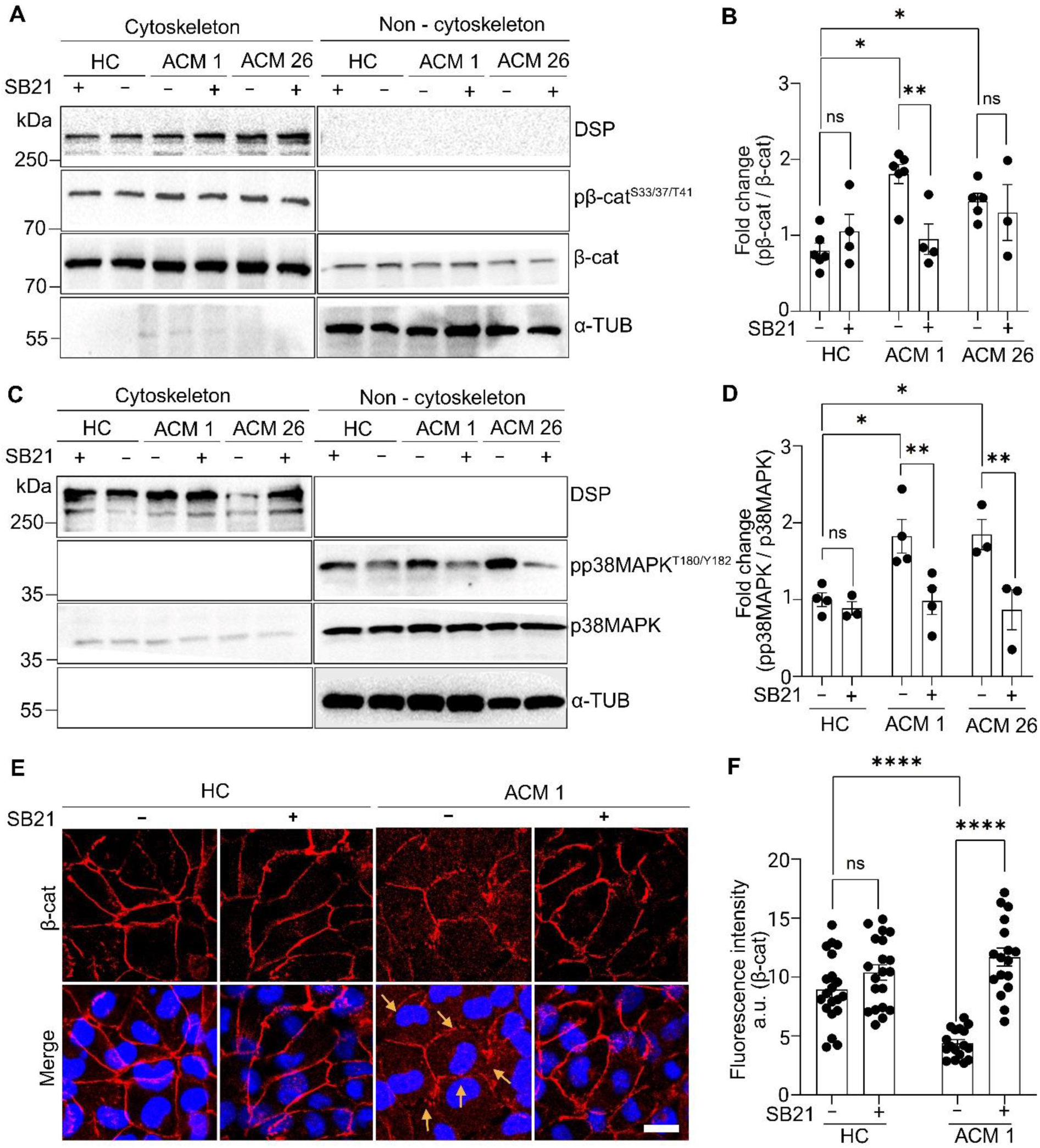
GSK-3β inhibition restores β-cat along cell junctions and inactivates p38MAPK (A-D) Triton X-100 analysis was performed in HL-1 cells for extraction of lysates after incubation with (+) and without (-) SB21 added 30 min prior to incubating with ACM 1, ACM 26 and HC-IgGs for 1 h, followed by Western blots and quantifications (B and D). To detect the protein expressions, fold change of ratio between respective phospho-protein and total protein was quantified (pβ-cat / β-cat), (pp38MAPK / p38MAPK) and normalized to HC-IgG. Data represent as mean ± SEM of 3 - 6 experimental repeats. **(E)** Immunofluorescence stainings in HL-1 cells to detect the β-cat abundance on the membrane were performed after 24 h pre- incubation of ACM 1 and HC- IgGs with (+) and without (-) SB21. **(F)** β-cat mean fluorescence intensity analysed by ImageJ. Each data points correspond to the 3 images from all conditions performed in 4 repeats. Data represents mean ± SEM. Statistical analysis was performed by two–way ANOVA with the Holm-Šidák multiple comparisons test. *p<0.05, **p<0.005, ***p<0.0005, ****p<0.00005 and ns-not significant.

**Figure 7.**
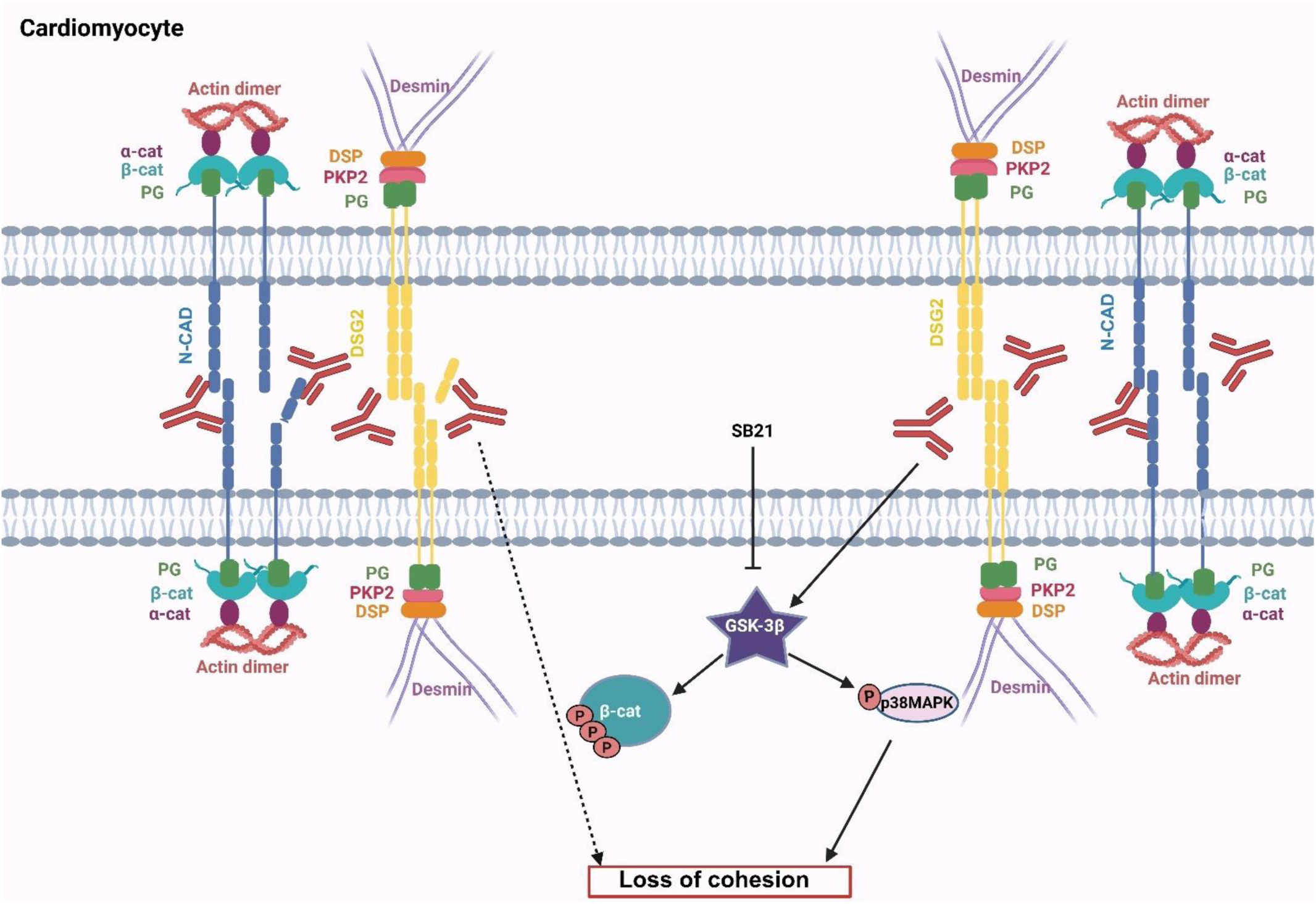
Mechanisms of ACM-IgG induced loss of cohesion in cardiomyocytes. ACM patients IgGs bind to DSG2 and activate GSK-3β leading to inactivation of β-catenin (β- cat) and activation of p38MAPK leading to loss of cardiomyocyte cohesion. On the other hand, ACM-IgGs contain catalytic activity leading to partial cleavage of DSG2 thereby partly responsible for cardiomyocyte cohesion loss. GSK-3β inhibition eventually ameliorates ACM-IgG induced cardiomyocyte cohesion loss.

## Discussion

ACM is a genetically inherited familial heart disease characterized by pathogenic genetic variants primarily found in genes encoding desmosomal and a few non-desmosomal proteins. Recent studies have established that inflammation plays an important role in ACM pathogenesis (28-30). Moreover, autoantibodies against ICD proteins and DSG2 have been identified in ACM patients in studies from three different laboratories (16, 17, 24). In the current study, we demonstrated that ACM-IgG from a few patients reduces cardiomyocyte cohesion in both human ACM hiPSC-CMs and in murine cardiomyocytes and that patient autoantibodies colocalize with DSG2 in living cardiomyocytes and impair DSG2 binding at the molecular level. Moreover, we have identified that GSK-3β negatively regulates cardiomyocyte cohesion and is activated by autoantibodies. This caused p38MAPK activation as well as phosphorylation of β-cat, which was paralleled by loss of β-cat from cell junctions. Taken together, these data support the hypothesis that autoantibodies could contribute to ACM pathogenesis by GSK-3β-mediated loss of cardiomyocyte cohesion.

First, we utilized IgGs from two ACM patients from our previous study, each carrying different pathogenic variants in *DSP* (ACM 1) and *PKP2* (ACM 15) (24). In hiPSC-CMs, both ACM-IgG fractions significantly reduced cohesion and co-localized with DSG2. Additionally, both ACM-IgGs co-localized with DSG2 in HL-1 cells. We collected new IgG samples from ACM patients with pathogenic variants *DSP* (ACM 28), *PKP2* (ACM 22, 26, and 27), and *DSG2* (ACM 29), as well as from 3 HR of ACM 1. Dissociation assays in HL-1 cells revealed that HR-IgGs do not cause a loss of cohesion. However, among the ACM-IgGs tested, three out of five fractions (ACM 22, 26, and 27-IgG) caused a significant increase in the number of fragments, albeit with varying degrees. We performed immunostaining of human ventricular cardiac tissue to check for the presence of anti-ICD IgGs in this cohort of ACM patients. To our surprise, all tested HR and ACM-IgG fractions exhibited staining of the ICD. These data indicate that both pathogenic and non-pathogenic autoantibodies are found in ACM patients, whereas in HR, autoantibodies are non-pathogenic. This may be explained by a process in which the first formation of non-pathogenic autoantibodies occurs independently of pathogenic variants in desmosomal proteins. During the course of the disease, in some patients, autoantibodies targeting the extracellular domain of DSG2, which is critical for cardiomyocyte cohesion are developed (31). A similar phenomenon is known from pemphigus patients and referred to as antigen-spreading (32-34). However, the observation that autoantibodies against ICD components are present in healthy relatives of patients but not in control individuals suggests that familial clustering is caused by genetic predisposition. In this line of thought, autoantibodies are not formed secondary to cardiomyocyte injury but most likely are present already in the concealed phase of the disease and thus before diagnosis. The presence of autoantibodies may potentially serve as a diagnostic criterion for ACM patients who are gene elusive.

Moreover, the results demonstrate that detection of autoantibodies against ICD proteins by immunostaining is very sensitive but does not reveal whether autoantibodies are pathogenic, which would require functional tests, including dissociation assays, cleavage experiments, or AFM studies, since all these assays require standardization, they cannot be applied in routine diagnostics.

We also isolated IgGs from two murine ACM models, *Pkp2^-/-^*and *Jup^-/-^*, respectively, at time points where the ACM phenotype was overt (21). Dissociation assays using IgGs from *Pkp2^-/-^*and *Jup^-/-^* mice did not reveal any loss of cohesion in HL-1 cells, similar to the IgGs from their corresponding wild-type littermates. Immunostaining with IgG fractions derived from these *Pkp2^+/+^, Pkp2^-/-^* model also did not display any positive staining of ICD. These results highlight that the development of autoantibodies against ICD proteins or DSG2 occurs through a different mechanism in ACM patients compared to murine models, which cannot be used to study the mechanisms underlying autoantibody formation. Indeed, it is documented that significant differences exist in innate and adaptive immunity of mice and humans, including immune system development and activation (35).

Thus, given that ACM is a desmosome-related disease which shares some immunological features with pemphigus, the fact that autoantibodies reduce cell cohesion in cultured murine cardiomyocytes, in murine slice cultures and also in patient-derived hiPSC-CMs is sufficient to conclude that loss of cell cohesion most likely contributes to pathogenesis of the disease (36). However, in patients, it is unknown to what extent and in which phase of the disease, loss of cell cohesion and thus autoantibodies may modulate disease progression. Since autoantibodies are absent in ACM mouse models, this could be tested in murine models only after passive transfer of autoantibodies, which may require repetitive applications over long time periods.

We then tested the catalytic properties of ACM-IgGs that caused a significant loss of cohesion. Both ACM 26 and ACM 27-IgG resulted in the cleavage of the DSG2 extracellular fragment, which was reduced in the presence of protease inhibitors, while cleavage was not evident in samples treated with ACM 22-IgG. Furthermore, we investigated whether ACM-IgGs induce a loss of DSG2 binding on the single molecule level. To address this, we conducted AFM experiments and measured DSG2 binding in the presence of ACM 22, ACM 26, and ACM 27-IgGs under cell-free conditions. All three ACM-IgGs significantly decreased the binding frequency of DSG2. Interestingly, ACM 22-IgG reduced DSG2 binding at the molecular level, as revealed by AFM, and reduced cardiomyocyte cohesion in dispase assay without causing DSG2 cleavage. Which suggests that ACM 22-IgG reduced DSG2 interaction most likely by direct inhibition, referred to as steric hindrance. Indeed, in the desmosome-related skin disease pemphigus vulgaris, it is well established that pathogenic autoantibodies against DSG3 cause loss of intercellular cohesion by directly interfering with desmosomal cadherin trans-interactions (37) (38).

We further investigated the signaling pathways activated by ACM-IgGs. In our previous study, we demonstrated that some ACM-IgGs activate p38MAPK (24). In this study, we focused on GSK-3β signaling, as this pathway was shown to be active in ACM patients, and its inhibition reversed the pathological features of ACM in mouse models and hiPSC-CMs from ACM patients (11, 26, 27, 39, 40). Similarly, GSK-3β was also found to be activated by autoantibodies of pemphigus patients (41).

We observed that under resting conditions, the inhibition of GSK-3β enhanced cardiomyocyte cohesion both in cultured murine cardiomyocytes and in murine myocardial slice cultures, indicating that GSK-3β is involved in the regulation of cell cohesion in cardiomyocytes. Moreover, in the *Jup^-/-^*murine ACM model, GSK-3β inhibition enhanced cardiomyocyte cohesion, revealing that inhibiting GSK-3β could strengthen desmosomal adhesion in pathophysiological conditions such as ACM. Similarly, when ACM-IgGs were applied, we observed that ACM 1, ACM 26 and ACM 27-IgG but not ACM 22-IgG significantly activated GSK-3β. Furthermore, GSK-3β-mediated phosphorylation of β-cat (S33/37/T41) was evident in cells incubated with ACM 1, ACM 22, and ACM 26-IgG. We focused on β-cat phosphorylation as proof of principle to demonstrate the downstream function of GSK-3β, leading to the degradation of β-cat and suppressing the WNT signaling pathway (4, 25, 39, 42). Indeed, ACM 1-IgG reduced β-cat at the cell junctions of HL-1 cell borders suggesting β-catenin degradation. Similarly, p38MAPK was activated by ACM1, ACM22 and ACM26-IgG. Since GSK-3β inhibition blunted both β-cat phosphorylation and p38MAPK activation, these data support that GSK-3β is involved in these events. Several studies have indicated that p38MAPK can be both downstream or upstream of GSK-3β (43, 44). In line with this, GSK-3β inhibition restored ACM-IgG-mediated loss of cohesion in HL-1 cells.

Since autoantibodies are absent in murine ACM models but present in human ACM patients, we further investigated the role of autoantibodies in a human ACM model. To do so, we isolated hiPSCs from the ACM 1 patient family, including hiPSCs from the healthy spouse of the ACM 1 patient, and also utilized her IgGs (HR 1) in this study. As an ACM model, we isolated hiPSCs from the daughter of the ACM 1 patient, who died of SCD and was post-mortem diagnosed as an ACM patient (22). ACM 1-IgG caused a significant loss of cohesion in both healthy and ACM hiPSC-CMs, whereas ACM 22-IgG was pathogenic in ACM hiPSC-CMs only. Similar to murine cardiomyocytes, GSK-3β inhibition completely abolished the loss of cohesion induced by ACM 1-IgG in healthy and ACM hiPSC-CMs. These results align with previously published studies where inhibition of GSK-3β rescued murine models from the ACM phenotype (11, 26, 40). Although our data from murine ACM models indicate that in these models GSK-3β is activated by the pathogenic variant and not via autoantibodies, however, in patients, GSK-3β activation may also be caused by autoantibodies.

In summary, our study gives important insights into the role of pathogenic and nonpathogenic autoantibodies in ACM and healthy patient relatives and supports the hypothesis that GSK-3β is important for ACM pathogenesis. Moreover, immunostaining can be used to detect ICD autoantibodies. Because several different methods, including co-staining in living human and murine cardiomyocytes, ELISA and also functional experiments such as dissociation and cleavage assays and AFM interaction studies all demonstrate that at least some autoantibodies target DSG2 and impair DSG2-mediated cohesion, it is unlikely that autoantibodies are not relevant in ACM but rather are an unspecific phenomenon.

## Materials and methods

### Reagents

The supplementary file details all the important reagents, mediators and antibodies used in this study and the missing methods in the main manuscript.

### Human blood collection and IgG isolation

Blood samples were collected from six healthy controls (HC), three healthy relatives (HR), seven ACM patients, two different murine ACM models (*Pkp2^-/-^* and *Jup^-/-^*) and their respective wild-type littermates (*Pkp2^+/+^* and *Jup^+/+^*) centrifuged to separate the sera, further the IgG fractions were separated by using protein A affinity chromatography as described previously (24, 45).

### HL-1 cell culture

HL-1 cells, immortalized murine atrial cardiomyocytes, were maintained in the Claycomb medium (cat #51800C) supplemented with fetal bovine serum (FBS, cat #F2442) and norepinephrine (NE) incubated at 37 °C and 5% CO2 in 0.02% gelatin and 25 µg/ml fibronectin-coated T25 flasks (46). Following 100% confluency, cells were passaged using trypsin-EDTA for dissociation. The cells were seeded at different densities for respective experiments and were supplemented with Claycomb medium devoid of norepinephrine (24, 47).

### Murine ACM models blood collections and IgG isolation

Mice of both genders were used for experiments at 12-weeks of age. Wild-type murine cardiac slices used for immunostainings were collected as described previously (21). For whole blood collection from mice, syringes and needles were pre-coated with 0.5M EDTA to prevent clotting during collection. Whole blood was collected into EDTA-containing micro sample tubes (1.6 mg/ml blood; Sarstedt, cat #41.1504.005) immediately after euthanization and thoracotomy by puncture of the right ventricle using a 23-25G needle, and centrifuged for 400 g x 5 min at 4°C. Supernatant was transferred to a new micro sample tube and stored at -80 °C until further use. IgG fractions were separated by using protein A affinity chromatography as described previously (24, 45).

### Human Ventricular cardiac tissue

The postmortem human ventricular cardiac tissue was obtained, and the fixation was carried out as described in the protocols mentioned in previous studies (24).Subsequently, the tissue was sectioned into 7 µm-thick slices using the cryostat (Thermo Fischer Scientific™, Cryostar NX70) and mounted onto the adhesive-coated glass slide to perform immunostaining with respective IgGs was performed as described previously (24).

### Human induced pluripotent stem cells (hiPSCs)

HiPSC lines used in this study were named as NP0040-8 [healthy male] (48), NP0151 [ACM (DSP. pGlu 952*) female], and NP0147 [healthy female] cell lines (22). Isolation of hiPSCs and characterization was done as described (22).

### Maintenance of hiPSC lines

HiPSC lines were cultured in complete Essential 8™ media in Vitronectin or Matrigel-coated 6-well plates in an incubator at 37 °C and 5% CO2 until they reached 70-80% confluency. Subsequently, cells were passaged by washing twice in DPBS without calcium and magnesium (Gibco™, cat #14190250), then dissociating in Versene for 3-5 min at room temperature. Versene was removed, and the detached cell colonies were resuspended in complete Essential 8™ medium with 2 µM ROCK inhibitor Y27632 (Adooq, cat #A11001-5) (NP0040-8 cell line was suspended in Essential 8™ medium without ROCK inhibitor). The cell cluster (8-10 cells) was divided at a ratio between 1:6 and 1:20 in either Matrigel® or Vitronectin-coated 6-well plates. Medium was changed daily with fresh complete Essential 8™ medium.

### Differentiation of hiPSC lines

HiPSCs were seeded as single cells to allow for differentiation into cardiomyocytes. HiPSC at optimum confluency were washed twice in DPBS without magnesium and calcium, then incubated for 5 min at 37 °C in TrypLE^TM^ Express (Gibco™, cat #12604021) with 10 µl DNase (Thermo Fischer Scientific^TM^, cat #90083) per 1 ml TrypLE^TM^. Complete Essential 8™ medium with 10 µM ROCK inhibitor Y27632 (Adooq, cat #A11001-5) was added to stop the TrypLE™ reaction and cells were then passed through a 40 µm cell strainer (corning^®^, cat #431750) to obtain single cells, followed by centrifugation at 200 g x 5 min. Cell pellets were resuspended in fresh complete Essential 8™ Medium containing 10 µM ROCK inhibitor. Single cells ranging from 0.4 - 0.8 x 10^6^ were seeded into wells of a Matrigel^®^-coated 6-well plate. Cells were supplemented with fresh complete Essential 8™ medium until they reached 80% confluency. On day 0, the cells were treated with 8 µM CHIR99021 (Sigma-Aldrich, cat #SML1046) for 24 h in RPMI GlutaMAX^TM^ (Gibco™, cat #61870010) medium supplemented with 50 µg/ml ascorbic acid (WAKO Chemicals Europe, cat #013-12061), B27 supplement without insulin (Thermo Fischer Scientific™, cat #A1895601) and penicillin/streptomycin (Gibco™, cat #15140-122). On day 1, the medium was replaced with new RPMI-B27 without insulin and cultured for 48 h. On day 3, cells were treated with 5 µM XAV939 (Sigma Aldrich, cat #X3004-5MG) and 5 µM IWP2 (Tocris, cat #3533/10) and incubated for 48 h in the reagents. On day 5, medium was replaced with fresh RPMI-B27 without insulin, followed by fresh RPMI-B27 with insulin on day 7. Further, the medium was changed every other day using RPMI-B27+Insulin, until the cells were used for cardiomyocyte purification.

### Replating and cardiomyocyte purification

Differentiated cardiomyocytes were replated in cardiomyocyte purification medium between days 8 and 14 after CHIR99021 treatment. The cells were rinsed twice with DPBS free of magnesium and calcium, followed by incubation with TrypLE™ Express containing DNase for 30 min. The reaction was stopped by adding RPMI-B27 with insulin, which was supplemented with 10 µM ROCK inhibitor. The cells were pelleted at 200 g x 5 min, and resuspended in fresh RPMI-B27 with insulin, including ROCK inhibitor, followed by seeding of 1 - 2 x 10^6^ cells per well on a Matrigel-coated 6-well plate. After 24 h, the medium was changed to RPMI-B27 with insulin and then to cardiomyocyte purification medium RPMI without glucose (Thermo Fischer Scientific™ cat #53003018) in 50 µg/µl ascorbic acid, L-Lactate, B27 with insulin, and penicillin/streptomycin for 3 days. The pure cardiomyocytes were then replated for further experiments.

### Dispase-based dissociation assays in HL-1 cells and hiPSC-CMs

A 24-well plate coated with gelatin and fibronectin was used to seed 0.15 x 10^6^ HL-1 cells, whereas a Matrigel^®^-coated 24-well plate was used to seed 0.4 x 10^6^ hiPSC-CMs. The HL-1 cells were cultured for 72 h, hiPSC-CMs underwent medium changes of RPMI-B27 with insulin every other day and were left for 18–20 days after replating. These cells were then incubated with IgGs for 24 h. For dissociation assay, the cells were washed twice with 500 µl HBSS (Sigma, cat #H1387-10L), Liberase-DH (0.065 U/ml, Sigma – Aldrich, cat #5401054001) and Dispase II (2.5 U/ml, Sigma-Aldrich, cat #D4693-1G) were then added and incubated at 37 °C until the cell monlayer detached from the well’s surface. Dispase II and Liberase-DH were carefully swapped out from HBSS. Applying the proper mechanical stress of 10–20 min for HL-1 cells and 1–10 min for hiPSC-CMs was done using an orbital shaker (Stuart SSM5 orbital shaker). Using a binocular stereomicroscope (Leica Microsystems, Wetzlar, Germany), images of wells with cell fragments were taken, and cell fragments were counted using ImageJ.

### Immunostaining in HL-1 cells, hiPSC-CMs and cardiac ventricular tissue slices with IgGs

Immunostaining in HL-1 cells and cardiac ventricular slices was performed as mentioned in previous studies (24, 47). In brief, HL-1 cells were seeded at a cell density of 0.15 x 10^6^ on 12 mm coverslips in a 24-well plate and grown for 72 h before incubating with the respective IgGs. Snap frozen human and murine ventricular slices were fixed with 2% Paraformaldehyde (PFA) and utilized further for immunostaining. Prior to fixation, HL-1 cells were treated with IgGs for 24 h and 1 h. Ventricular tissues were incubated with the respective IgGs post-PFA fixation. For immunostainings, cells and tissue were fixed with 2% paraformaldehyde (PFA), then washed with 1X PBS and permeabilized with 0.1% Triton X-100 (for cells) / 1% Triton X-100 (for cardiac tissue). Blocking the cells and tissue was done by Bovine serum albumin/ normal goat serum (BSA/NGS) in a wet chamber. The respective primary antibodies for HL-1 cells and IgGs from human sera for human cardiac slices, murine wild-type and *Pkp2^-/-^ A*CM-IgGs for murine cardiac slices were incubated overnight at 4 °C, washed 3X times each 5 min, and then fluorophore-coupled secondary antibody of the same species was incubated for 1 h at room temperature (RT) followed by 3X times washes with PBS. For detecting primary antibodies goat anti-human, goat anti-mouse and goat anti-rabbit secondary antibodies were used. Mounting was done using Pro-Long Diamond antifade mounting medium (Thermo Fischer Scientific^TM^, cat #P36961).

HiPSC-CMs were seeded at 0.2 X 10^6^ per Matrigel^®^-coated ibidi chambers (Ibidi LOT #230725/4) and the experiments were performed between days 20–24 after CHIR99012 treatment. After treatment of cells with respective IgGs for 1 h, the cells were fixed using 2% PFA, followed by permeabilization with 0.25% Triton X-100 in 0.1 M PBS. The blocking was done in BSA/NGS followed by primary antibody incubation at 4° C overnight, subsequently washing with 1x PBS for 5 times and 1 h incubation of fluorophore coupled secondary antibody of same species at RT, mounting was done using DABCO (1,4 diazabicyclo 2,2,2 octane, DABCO^®^, Carl Roth, cat #0718.2). Leica SP5II confocal software with an 63x oil objective and LAS-AF software were used for the image acquisition and analysis.

### Immunostaining for Stimulated emission depletion (STED) microscopy

HL-1 cells were seeded at 0.15 x 10^6^ on 12 mm cover slips, as stated in the methods sections. Following fixation with 2% PFA and permeabilization with 0.1% Triton-X-100 in PBS. Afterward, the samples were washed 3X with 50 mM ammonium chloride. Blocking was performed using BSA/NGS for 60 min at RT, followed by overnight incubation at 4 °C with the primary antibody. The next day, samples were washed 3X with 50 mM ammonium chloride in PBS, then incubated for 1 h at RT with a species-specific, STED-compatible fluorophore-conjugated secondary antibody. Mounting was done using Pro-Long Diamond antifade mounting medium (Thermo Fischer Scientific^TM^, cat #P36961). Images from a STED microscope equipped with an Abberior Expert line setup with a 100x oil objective were analyzed using Imspector image acquisition software (Abberior Instruments GmbH, Göttingen, Germany) (49). The STED effect was achieved with 75 nm range, 775 nm pulsed laser, 20 nm pixel focus, 50% laser power, 80% resolution.

### Quantification of IgG colocalization with DSG2

ImageJ program was used to quantify the colocalization of IgG with DSG2 (Fiji: ImageJ-win64). The dialog box coloc 2 from the colocalization analysis was selected after converting the images from the channels of interest to 8-bit type. The necessary algorithms imply the use of the Pearson correlation coefficient. For colocalization, the mean of the Pearson’s r-value was considered where 0 is no correlation, between 0 to1 is positive correlation and between 0 to - 1 is negative correlation.

### Enzyme-Linked Immunosorbent Assay (ELISA)

A 96-well plate (Maxisorp, Thermo Fisher Scientific) was coated with 50 µl/well of 2.5 µg/ml hDSG2-his (Medchem Express) in DPBS (Sigma Aldrich), pH 7.4 and incubated for 1 h at RT. Wells were blocked with 100 µl of blocking buffer (1% (w/v) BSA (Sigma Aldrich) and 0.05% (v/v) Tween20 (Sigma Aldrich) in DPBS). Blocked wells were washed 3X with PBST (0.05% (v/v) Tween20 in DPBS) and incubated with 1:5 serial dilutions of donor IgG samples (concentrations: 500 nM/ 75 µg/ml; 100 nM/ 15 µg/ml, 20 nM/ 3 µg/ml) and one blank sample only containing PBS for 1 h at RT. After 3 washes with PBST, a polyclonal murine anti-human Fc primary antibody (3 µg/ml in blocking buffer, kindly provided by H. Flaswinkel, E. Kremmer and H. Leonhardt) was added and incubated for 1 h at RT. Wells were washed 3X and secondary antibody was added (1:10000, goat anti-mouse-HRP; Jackson Immuno Research, cat #AB_2338506). After 1 h at RT, wells were washed 3X with PBST and developed by adding 50 µl of TMB substrate (ScyTek Laboratories). Absorbance at 605 nm wavelength was measured using a plate reader (Tecan Spark) and values were background-subtracted [A (cIgG=x) - A (cIgG=0)].

### DSG2-Fc based Cleavage assay

Cleavage assay was performed utilizing recombinant human extracellular domain of DSG2 tagged with Fc (DSG2-Fc), as described in our previous publication (20) but with some modifications. The ACM-IgGs and HC-IgG were incubated for 10 min at RT with cleavage buffer (10 mM HEPES, pH 7.5; 0.02% NaN3; 0.05% Brij 35) containing DMSO (1% v/v), with and without protease inhibitor (cOmplete™: cat #11697498001; cOmplete™, EDTA-free: cat #11873580001, Roche). Following that, 200 ng of hDSG2-Fc (DSG2-512H, cat #CAA81226, Creative BioMart) was added to the 10 µl final reaction mixture and incubated at 37 °C for 24 h. To detect the DSG2-Fc protein, Western blot was performed using specific primary antibody against DSG2 (Origene, cat #BM5016) and horseradish peroxidase-linked goat anti-mouse secondary antibody.

### Atomic force microscopy (AFM)

Atomic force microscopy measurements were performed as described before (24). Briefly, Nanowizard® III AFM (JPK Instruments, Berlin, Germany) with an optical fluorescence microscope (Axio D1 observer Carl Zeiss) was used at 37 °C. AFM cantilevers (MLCT AFM Probes, Bruker, USA) were coated with recombinant DSG2-Fc as previously described (50). Force-Distance curves (FDC) recorded with a 0.3 nN loading force, 0.1 s contact delay, and 1 µm/s retraction velocity. Each experiment recorded 1000 FDC in HBSS buffer to establish baseline trans-interaction probability. An additional 1000 FDCs were recorded after 1 h incubation of mica and cantilevers with HC and ACM-IgGs. JPKSPM data processing software was used to analyze FDC unbinding signatures to determine interaction probabilities, normalized to baseline values to account for coating variations. Adhesion maps were generated by overlaying the position of occurring unbinding events with the mica topography maps.

### Triton X-100 extraction assay and Western blot

Following treatments, the HL-1 cells and hiPSC-CMs were incubated on ice in Triton extraction buffer (0.5% Triton X-100, 50 mM MES, 25 mM EGTA, 5 mM MgCl_2_, pH 6.8) supplemented with protease inhibitor (cOmplete protease inhibitor cocktail, Roche, CO-RO, cat #11836145001) and phosphatase inhibitor (PhosStop, Roche, PHOSS-RO, cat #4906837001). The cells were scraped using a cell scraper and collected in an Eppendorf tube, then spun at 15000 g for 5 min at 4 °C in a centrifuge (Eppendorf 5430R) to separate cytoskeleton from non-cytoskeleton proteins. The non-cytoskeleton fractions were collected in a separate Eppendorf tube, while the cytoskeleton fractions were lysed in SDS lysis buffer (12.5 mM HEPES, 1 mM EDTA disodium, 12.5 mM NaF, and 0.5% SDS, pH 7.6) containing protease and phosphatase inhibitors and sonicated (pulse rate 10 times). Protein concentrations were quantified using a BCA assay kit (Pierce BCA assay kit cat #A55864, Thermo Fischer Scientific™) and the sample was prepared with Laemmli loading buffer (including 50 mM DTT). Furthermore, a 10% polyacrylamide gel (PAGE) was used for the Western blot, which was then transferred to a nitrocellulose membrane. Blocking for 90 min was performed with either 5% Milk/TBST or 5% BSA/TBSTor 1X Rotiblock, followed by an overnight incubation at 4 °C with the primary antibody, washed and 1 h incubation at RT with the same species horseradish peroxidase-linked secondary antibody. The membrane was developed using Image developer (iBright 1500, Thermo Fischer Scientific™).

### Ethics approval

Animal handling, housing and husbandry were performed in accordance with the guidelines from the Directive 2010/63/EU of the European Parliament and were approved by the regional government of Upper Bavaria (Gz. ROB-55.2-2532.Vet_02-19-172, for Jup mice) or the local ethics committee of the government of Lower Franconia (RUF-55.2.2-2532-2-663 and 55.2.22-2532-2-955 for Pkp2 mice).

Blood sample collection and IgG isolation from ACM and healthy individuals was performed with the approval of Ethics Committee at LMU (20-1061) and informed written consent was obtained.

Human heart ventricular tissues were collected from body donors at the Institute of Anatomy and Cell Biology, LMU, Munich, Germany. Written informed consent was obtained from the female body donor for the use of tissue samples in research. The body arrived within 17 h after decease and was free of any cardiac disorders.

The Ethics Committee of the Medical Faculty of the University of Cologne (authorization number 14-306) approved patient tissue collection and use for generating hiPSCs. Further hiPSC study at the LMU was approved by the Ethics Committee approval at LMU (23-0203).

### Statistical analysis

GraphPad Prism version 10 software (GraphPad Software, La Jolla, USA) was used for statistical analysis, which included un-paired t-tests, one and two-way ANOVA, p-values < 0.05 were considered significant.

## Author contributions

SP, KS, EK, SM, MH and SY acquired and analyzed the data. MR and AS established and analyzed DSG2-ELISA. TW and BG provided IgG samples from *Pkp2^+/^*^+^ and *Pkp2^-/-^* mice. RB, SC and ST were involved in collecting the blood samples. hiPSCs were generated, and protocols to differentiate hiPSCs into cardiomyocytes were established in the lab of TŠ. SP and SY drafted the manuscript. SY, and JY designed the research and made critical revisions to the manuscript for important intellectual content. All authors proofread the manuscript.

## Acknowledgements

We thank Kilian Skowranek for his technical assistance in isolating the IgGs and Stefanie Jeschke for mice handling and Prof. Heinrich Leonhardt for critical discussion and infrastructure support for the ELISA results.

