## Supplementary file for "Autoantibodies in patients with arrhythmogenic cardiomyopathy activate GSK-3β resulting in a loss of cardiomyocyte cohesion"

### Supplementary methods

#### Western blotting

A 10% polyacrylamide stacking and resolving gel was prepared for SDS-PAGE. Electrophoresis was carried out using the Bio-Rad Mini-PROTEAN® Tetra Cell system, initiated at 80 V for stacking, followed by 120 V for 60 min to resolve the proteins. Protein samples were mixed with Laemmli sample buffer containing dithiothreitol (DTT) and denatured by boiling at 95°C for 5 min. As a molecular weight reference, 5 µL of PageRuler™ Plus Prestained Protein Ladder (Thermo Fisher Scientific™, Cat. #26619) was loaded alongside the samples.

Following electrophoresis, proteins were transferred onto a 0.45 µm nitrocellulose membrane (Thermo Fisher Scientific™, Cat. #LC2006) at a constant current of 350 mA for 90 min. Membranes were then incubated for 10 min with the No-Stain™ Protein Labeling Reagent (Thermo Fisher Scientific™, Cat. #A44449) and imaged using the iBright 1500 imaging system (Thermo Fisher Scientific™).

To block non-specific binding sites, membranes were incubated at room temperature for 1 h in 0.1 M Tris-buffered saline containing 0.1% Tween 20 (TBST), supplemented with 5% non-fat dry milk (Sigma Aldrich, Cat. #70166-500G), 5% bovine serum albumin (BSA), or 1X Roti®-Block solution (Carl Roth, Cat. #A151.2).

#### Enzyme-Linked Immunosorent Assay

DSG2-Fc ELISA was performed following the same procedure as DSG2-His, using 5 µg/ml hDSG2-hFc (DSG2-512H, cat #CAA81226, Creative BioMart) for coating and IgG concentrations of 1500 nM(225 µg/ml), 500 nM(75 µg/ml) and 100 nM(15 µg/ml), performed in singlets. Detection of IgG bound to hDSG2-hFc was achieved using goat anti-Fab antibody (BioRad, cat #STAR159) and anti-goat HRP (Abcam, cat #ab97110). Blocking buffer and antibody solutions used for hDSG2-hFc ELISA were supplemented with 1 tablet per 50 ml of protease inhibitor cocktail (c0mplete, EDTA free; Roche). Absorbance at 450 nm wavelength was measured using a plate reader (Tecan Spark) and values were background-subtracted [A (cIgG=x) - A(cIgG=0)].

### Supplementary Figures

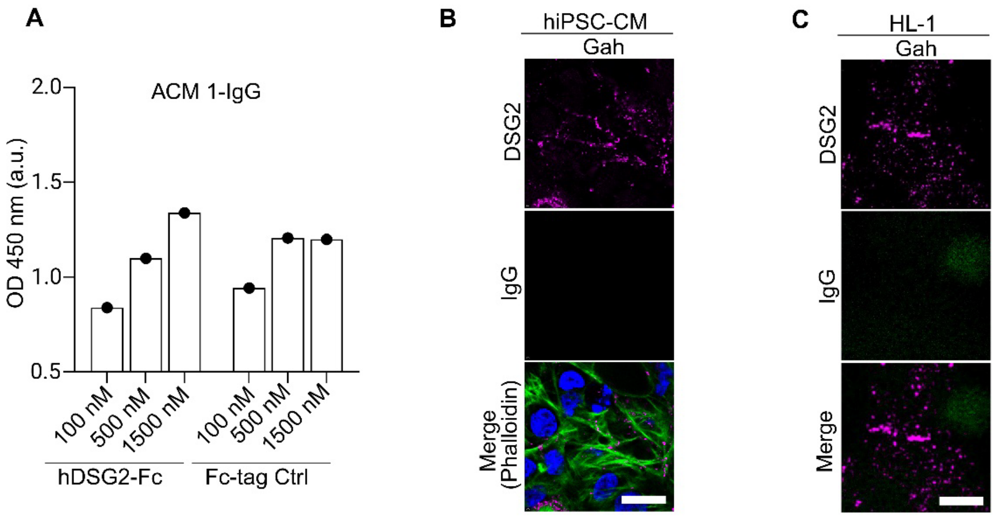

**Supplementary Figure 1. DSG2-Fc ELISA and secondary antibody control stainings.**

**(A)** ELISA with recombinant extracellular DSG2-Fc protein was used to detect the binding of patient-derived ACM 1-IgG to DSG2. 5 µg/ml hDSG2-hFc coated wells were incubated with 1:5 serial dilution of ACM 1-IgG at different concentrations (1500 nM, 500 nM and 100 nM). The absorbance (OD) was measured at 450 nm. The secondary antibody control stainings on **(B)** hiPSC-CMs and **(C)** HL-1 cells were performed using goat anti-human labelled Cy3. Phalloidin was used to stain sarcomeric actin as a hiPSC-CMs marker. Goat-anti-human (Gah)-IgG shows no cross-reactivity with DSG2. N-3, Scale bar 10 µm

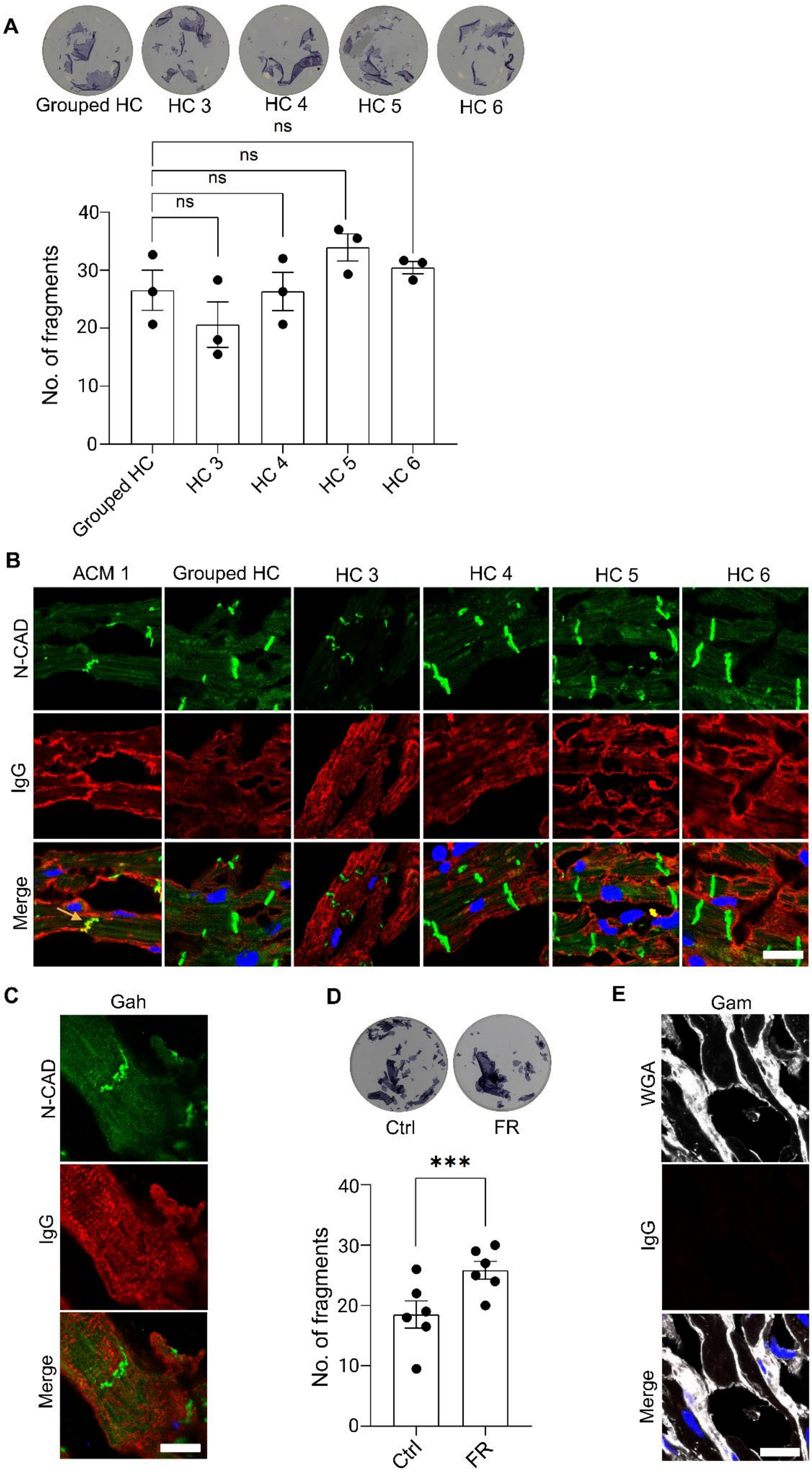
A
Grouped HC
HC 3
HC 4
HC 5
HC 6
ns
ns
ns
ns
No. of fragments
40
30
20
10
0
B
ACM 1
Grouped HC
HC 3
HC 4
HC 5
HC 6
N-CAD
IgG
Merge
C
Gah
N-CAD
IgG
Merge
D
Ctrl
FR
***
No. of fragments
40
30
20
10
0
Ctrl
FR
E
Gam
WGA
IgG
Merge

**Supplementary Figure 2. Dissociation assays and immunostaining in human and murine ventricular cardiac slices.**

**(A**) Dissociation assays performed using grouped control IgG (Grouped HC) and single control IgGs fractions (HC 3, HC 4, HC 5, HC 6) after 24 h treatment in HL-1cells. Representative images from 3 experimental repeats show cellular fragments in the top-panel. The bar graph shows mean ± SEM. One-way ANOVA with Holm-Šidák multiple comparisons test for statistical analysis. **(B)** Immunostaining analysis utilizing grouped control IgG and single control IgG was performed on human ventricular cardiac tissue. ACM 1-IgG was used as positive control. N-CAD used as an ICD marker. The single control IgGs do not show overlap with N-CAD. **(C)** The secondary goat anti-human (Gah) labelled with Cy3 used for detecting cross-reactive antigens in human ventricular tissue. Mild cross-reactivity with human antigens was observed, as evidenced by intracellular staining. **(D)** Dissociation assay using positive control FR (Froskolin/rolipram) after 1 h treatment in HL-1 cells shows enhanced cardiomyocyte cohesion compared to control (Untreated cells). The bar graph represents the mean ± SEM of 4 repeats. The un-paired t-test was performed for statistical analysis. **(E)**The goat anti-mouse antibody (Gam) labelled with Cy5 used as secondary antibody control. WGA used as membrane marker. Immunostaining revealed no cross-reactivity of Gam with murine cardiac tissue. *$p<0.05$ and ns-not significant.

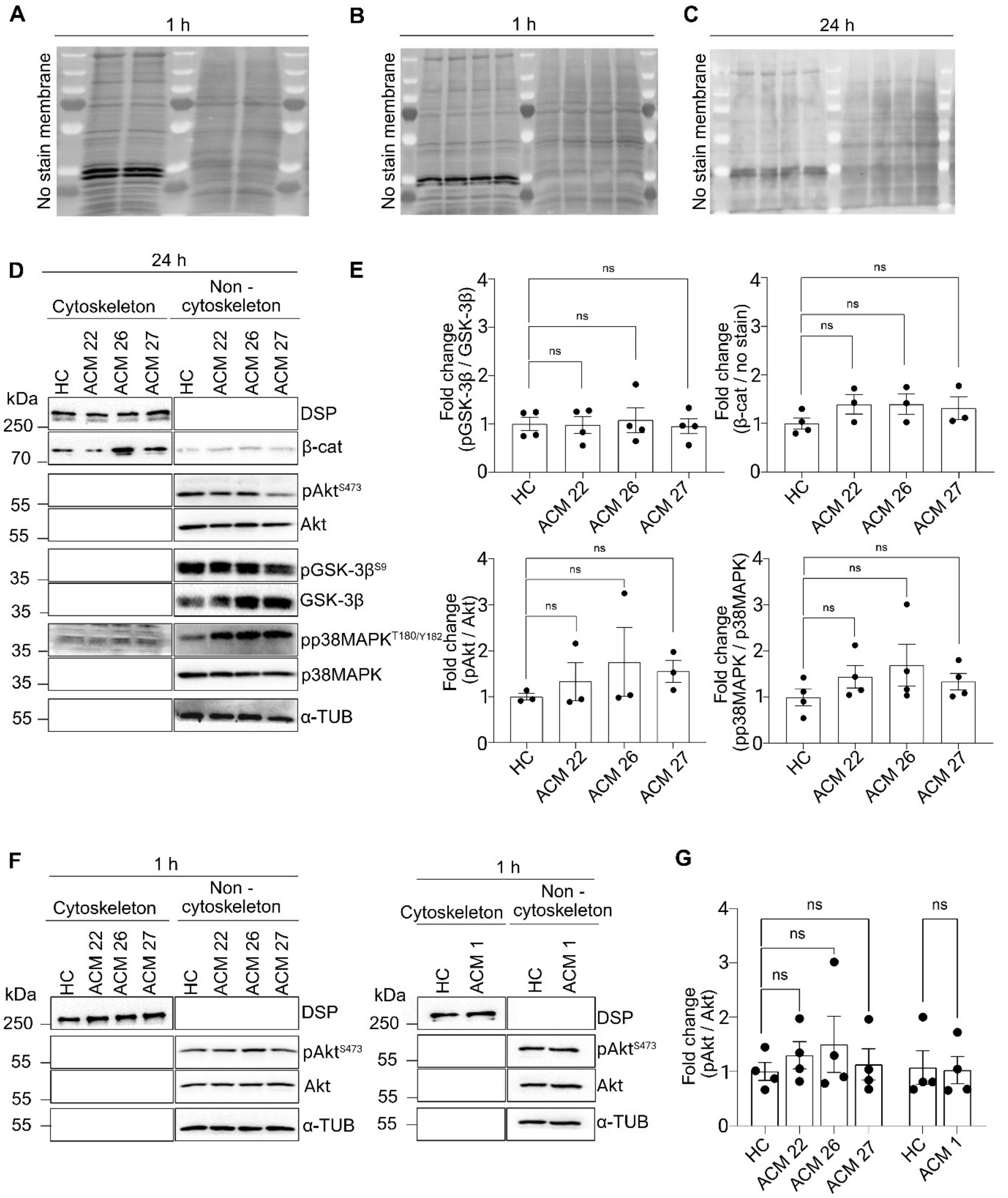

### Supplementary Figure 3. Triton-X-100 Assays

**(A - C)** The no-stain labelling buffer was used as a loading control in the Western blot. **(D and E)** The Western blot was performed after 24 h treatment of HC and ACM-IgGs for the expression analysis of respective proteins (GSK3-β, β-cat, pp38, Akt) in ACM-IgG normalized to HC-IgG. The bar graphs show the mean ± SEM of 3 - 4 repeats. **(F and G)** Triton X-100 with 1 h HC and ACM-IgGs treatments were carried out to detect the Akt protein expression. The bar graphs represent mean ± SEM of 4 experimental repeats. One-way ANOVA with

Holm-Šidák multiple comparison test and un-paired *t*-test were used for statistical analysis. *$p<0.05$ and ns-not significant.

**Table 1:** Arrhythmogenic cardiomyopathy (ACM) patient details

| Gene Mutation | Patient ID | Task force criteria | Age* | Sex | Clinics |
|---|---|---|---|---|---|
| DSP (c.2854G>T, p.E952*) | **ACM 1** | Yes | 63 | male | PPK, supraventricular arrhythmia, RV enlargement and + WMA, LVEF 50 % |
| PKP2 (c.369G>A) | **ACM 15†** | Yes | 68 | male | ventricular tachycardia, RV enlargement, severe RV dysfunction, RVEF 18 %, right ventricular assist device, LVEF 54 % |
| PKP2 (NM_004572.3(PKP2):c.2146-1G>C) | **ACM 22** | Yes | 18 | male | RV enlargement, reduced RV ejection fraction (15%), normal LVEF; LGE at RV, multiple slow VTs; RV thrombus; |
| PKP2 c.(1510+1_1511-1)_(1688+1_1689-1)del | **ACM 26** | No | 45 | female | Normal size of the RV, mild RV dyskinesia, normal RV-EF; normal LV-EF; frequent PVCs (burden 1%); |
| PKP2 c.(1510+1_1511-1)_(1688+1_1689-1)del | **ACM 27** | Yes | 22 | male | Normal size of the RV, normal RV-EF; normal LV-EF; history of sustained VT |
| DSP (c.2854G>T, p.E952*, Exon 20) | **ACM 28** | No | 42 | female | Subepicardial LGE LV posterior and lateral wall; LVEF 64%, RVEF 61%, no LV/RV dilatation; no arrhythmias, no syncope |
| DSG2 c.(1069_1072del), p.(Lys357GlnfsTer22) | **ACM 29** | Yes | 38 | female | Normal LVEF (60%), RV dilatation and reduced EF; multiple non-sustained VTs; history of resuscitation because of VF |

DSP: desmoplakin, LGE: late gadolineum enhancement, LV: left ventricle, LVEF: left ventricular ejection fraction, PKP2: plakophilin 2, PPK: palmoplantar keratoderma, RV: right ventricle, RVEF: right ventricular ejection fraction, WMA: wall motion abnormality. * Age at the time of blood collection

**Table 2:** Details of Antibodies

| **Antibody (species, Company, Catalog no.)** | **Antibodies dilution used for WB** | **Antibodies dilution used for IF** |
|---|---|---|
| DSP (mAb, mouse, Progen, #690003) | 1:1000 in Rotiblock | |
| DSG2 (mAb, mouse, Progen, #61002) | 1:1000 in BSA/TBST | |
| DSG2 (mAb, rabbit, Progen, #610121) | | 1:100 in BSA/NGS |
| DSG2 (mAb, mouse, Origene, #BM5016) | 1:200 in Rotiblock | |
| N-CAD (mAb, mouse, BD Transduction, #610921) | 1:1000 in BSA/TBST | 1:500 in BSA/NGS |
| GSK3-β (3D10, mAb, mouse, Cell Signaling Technology®, #9832) | 1:1000 in BSA/TBST | |
| pGSK3-β$^{Ser9}$ (mAb, rabbit, Cell Signaling Technology®, #5558) | 1:1000 in BSA/TBST | |
| pβ-catenin$^{Ser33/37/Thr41}$ (pAb, rabbit, Cell Signaling Technology®, #9561S) | 1:1000 in BSA/TBST | |

| | |
|---|---|
| β-catenin (mAb, mouse, BD Transduction, #610154) | 1:1000 in BSA/TBST |
| pp38MAPK$^{Thr180/Tyr182}$ (mAb, rabbit, Cell Signaling Technology®, #4511) | 1:1000 in BSA/TBST/ 1:1000 in Rotiblock |
| p38MAPK (pAb, rabbit, Cell Signaling Technology®, #9212) | 1:1000 in BSA/TBST/ 1:1000 in Rotiblock |
| pAkt$^{Ser473}$ (mAb, rabbit, Cell Signaling Technology®, #4060) | 1:1000 in BSA/TBST |
| Akt (pAb, rabbit, Celll Signaling Technology®, #9272 ) | 1:1000 in BSA/TBST |
| α-Tubulin (mAb, mouse, Abcam, #ab7291) | 1:4000 in BSA/TBST |
| Goat-anti-mouse-HRPO (Dianova, #115-035-068) | 1:10000 in TBST |

| | | |
|---|---|---|
| Goat-anti-rabbit-HRPO (Dianova, #111-035-045) | 1:10000 in TBST | |
| Goat anti-human Cy3 (Dianova, #109-165-008) | | 1:600 in PBS |
| Goat-anti-mouse Cy3 (Dianova, # 115-165-164) | | 1:600 in PBS |
| Goat-anti-rabbit Cy3 (Dianova, # 111-165-003) | | 1:600 in PBS |
| Goat-anti-mouse Cy5 (Dianova, # 115-175-071) | | 1:600 in PBS |
| Goat-anti-rabbit Cy5 (Dianova, # 111-175-144) | | 1:600 in PBS |
| Phalloidin Alexa 488 (Molecular Probes\Life technology, # A 12°9) | | 1:400 in PBS |
| Goat-Anti-rabbit STAR RED (Abberior, #STRED-1002) | | 1:200 in PBS |
| Goat-Anti-human Alexa 594 (Invitrogen, #A-11013) | | 1:200 in PBS |

| | | |
|---|---|---|
| DAPI, Roche, #10236276001 | | 1:2000 in PBS |
| Wheat germ agglutinin, WGA Alexa 488(Thermo Fisher Scientific, #W11261) | | 1:400 in PBS |

**Table 3:** Details of mediators

| Name | Concentration |
|---|---|
| Forskolin(F) (Sigma-Aldrich, #F3917) | 5 µM |
| Rolipram(R) (Sigma-Aldrich, #R6520) | 10 µM |
| SB216763 (MedChemExpress #HY12012) | 10 µM |
